# Generating single-cell gene expression profiles for high-resolution spatial transcriptomics based on cell boundary images

**DOI:** 10.1101/2023.12.25.573324

**Authors:** Bohan Zhang, Mei Li, Qiang Kang, Zhonghan Deng, Hua Qin, Kui Su, Xiuwen Feng, Lichuan Chen, Huanlin Liu, Shuangsang Fang, Yong Zhang, Yuxiang Li, Susanne Brix, Xun Xu

## Abstract

Stereo-seq is a cutting-edge technique for spatially resolved transcriptomics that combines subcellular resolution with centimeter-level field-of-view, serving as a technical foundation for analyzing large tissues at the single-cell level. Our previous work presents the first one-stop software that utilizes cell nuclei staining images and statistical methods to generate high-confidence single-cell spatial gene expression profiles for Stereo-seq data. With recent advancements in Stereo-seq technology, it is possible to acquire cell boundary information, such as cell membrane/wall staining images. To take advantage of this progress, we update our software to a new version, named STCellbin, which utilizes the cell nuclei staining images as a bridge to align cell membrane/wall staining images with spatial gene expression maps. By employing an advanced cell segmentation technique, accurate cell boundaries can be obtained, leading to more reliable single-cell spatial gene expression profiles. Experimental results verify that STCellbin can be applied on the mouse liver (cell membranes) and *Arabidopsis* seed (cell walls) datasets and outperforms other competitive methods. The improved capability of capturing single cell gene expression profiles by this update results in a deeper understanding of the contribution of single cell phenotypes to tissue biology.

**Availability & Implementation:** The source code of STCellbin is available at https://github.com/STOmics/STCellbin.

## STATEMENT OF NEED

Spatially resolved single cell transcriptomics enables the generation of comprehensive molecular maps that provide insights into the spatial distribution of molecules within the single cells that make up tissues. This groundbreaking technology offers insights into the location and function of cells in various tissues, enhancing our knowledge of organ development [1], tumor heterogeneity [2], cancer evolution [3], and other biological mechanisms. Resolution and field-of-view are two critical parameters in spatial transcriptomics. High resolution enables detailed molecular information at the single-cell level, and large field-of-view facilitates the creation of complete 3D maps that represent biological functions at the organ level. Stereo-seq simultaneously achieves subcellular resolution and a centimeter-level field-of-view, providing a technical foundation for obtaining comprehensive spatial gene expression profiles of whole tissues at single-cell level [4]. Our previous work offers the one-stop software StereoCell for acquiring high signal-to-noise ratio single-cell spatial gene expression profiles from Stereo-seq data [5]. StereoCell takes the cell nuclei staining image tiles and its corresponding spatial gene expression data as input, and its main process includes image stitching, image registration, tissue segmentation, cell nuclei segmentation and molecule labeling steps. The image data generated by Stereo-seq used for StereoCell are cell nuclei staining images. However, there is a big difference between cell nuclei and cell boundary staining images, based on cell membrane/wall staining, in terms of the ability to capture robust and precise cell specific gene expression profiles. Despite the widespread use of spatial techniques, such as MERFISH [6], CosMx [7], and Xenium [8], several of these techniques still struggle to achieve accurate cell boundary information, as they are based on cell nuclei staining images that can be generated using stains such as 4,6-diamidino-2-phenylindole (DAPI). Hematoxylin-eosin (H&E) and single strand DNA fluorescence (ssDNA) nuclei staining images are also commonly used and readily obtainable data. The updated Stereo-seq technology implement a procedure based on simultaneous cell membrane/wall and cell nuclei staining by adding multiplex immunofluorescence (mIF) and calcofluor white (CFW) staining [9,10], which can automatically acquire more accurate cell boundary information and thereby obtain more reliable single-cell spatial gene expression profiles.

We update StereoCell to a new version, named STCellbin. We retain the image stitching, tissue segmentation and molecule labeling steps from StereoCell and improve the image registration and cell segmentation steps. The “track line” is a crossed linear marker embedded on the Stereo-seq chip, which is the key information for the image registration step of StereoCell [5]. As the cell membrane/wall staining images miss the “track line” information, we utilize the cell nuclei staining images as a bridge to align the cell membrane/wall staining images with the spatial gene expression maps, upon which we obtain the registered cell boundary information in the cell segmentation step. Based on the cell boundary information, we directly assign the molecules to their corresponding cells, obtaining single-cell spatial gene expression profiles. We apply STCellbin on mouse liver (cell membrane) and *Arabidopsis* seed (cell wall) datasets and confirm the accuracy of cell segmentation. This update offers a comprehensive workflow to obtain more reliable single-cell spatial gene expression profiles based on cell membrane/wall information, providing support and guidance for related scientific investigations, particularly those based on Stereo-seq data.

## IMPLEMENTATION

### Overview of STCellbin

The process of STCellbin includes image stitching, image registration, cell segmentation and molecule labeling (Fig. 1). The Stereo-seq spatial gene expression data, cell nuclei and cell membrane/wall staining image tiles are input into STCellbin. The stitched cell nuclei and cell membrane/wall staining images are obtained through the MFWS algorithm [5]. These two stitched staining images are registered using the Fast Fourier Transform (FFT) algorithm [11]. The spatial gene expression data is transformed into a map, this map and a stitched cell nuclei staining image are registered based on “track lines”. Thus, the registration of the gene expression map and cell membrane/wall staining image is implemented. Cell segmentation is performed on the registered cell membrane/wall staining image by the adjusted Cellpose 2.0 [12] to obtain the cell mask. The molecules are assigned to their corresponding cells according to the cell mask to obtain the single-cell spatial gene expression profile as the output. The tissue segmentation step based on Bi-Directional ConvLSTM U-Net [13] is set as optional, which can generate a tissue mask to assist in filtering out impurities outside the tissue.

**Figure 1.**
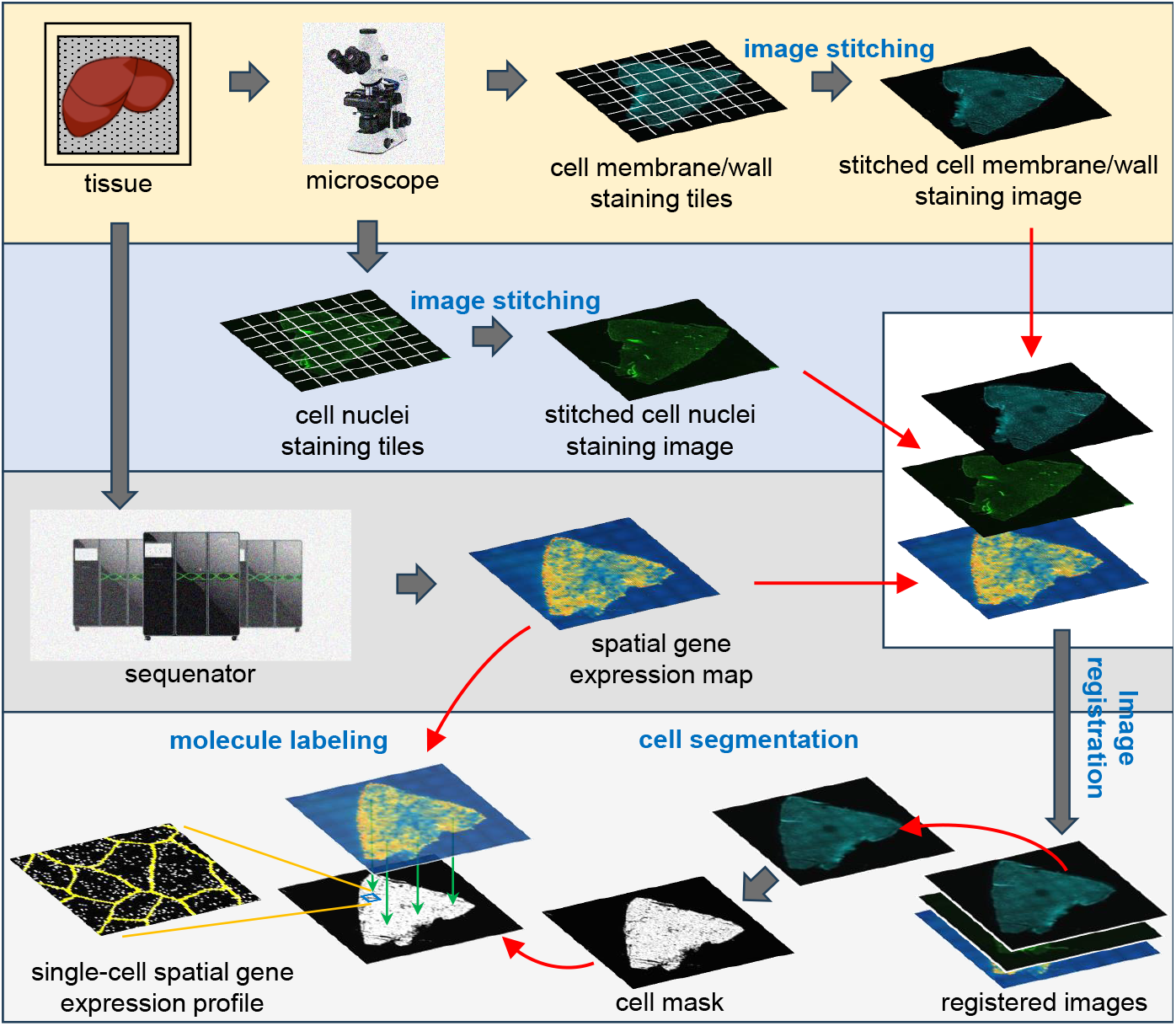
Overview of STCellbin. The cell nuclei and cell membrane/wall staining image tiles are stitched into individual large images respectively. The spatial gene expression map and stitched cell membrane/wall staining image are registered with the stitched cell nuclei staining image as a bridge. The cell mask is directly obtained from the registered cell membrane/wall staining image by cell segmentation. The single-cell spatial gene expression profile is obtained by overlaying the generated cell mask and the gene expression map.

### Image stitching

The image stitching step in STCellbin is consistent with it in StereoCell. The MFWS algorithm [5] is adopted, which calculates the offsets of two adjacent tiles with overlapping areas using FFT [11] to stitch these two tiles and extends this process to all tiles. The relative error, absolute error and computational efficiency of MFWS have been verified in our previous work [5].

### Image registration

The registration of STCellbin includes three stages. The first stage is the registration of the stitched cell nuclei and stitched cell membrane/wall staining images. These two staining images have similar sizes and no large difference in rotation because the chip does not move when they are photographed. The key to this registration is to calculate their offset. The size of the cell membrane/wall staining image is adjusted to be consistent with the cell nuclei staining image by cutting and zero-padding (Fig. 2A). The two staining images are mean-based subsampled [14] (Fig. 2B). The offset of the subsampled images is calculated through FFT [11], similar to MFWS [5] (Fig. 2C). The calculated offset is restored to the scale of the original images (Fig. 2D). Thus, these two staining images can be registered. The second stage is the registration of the stitched nuclei staining image and spatial gene expression map. This registration is the same as it in StereoCell [5]. The spatial gene expression data is transformed into a map. The stitched cell nuclei staining image is registered with the map based on “track lines” by performing scaling, rotating, flipping and translating on the stitched cell nuclei staining image. The third stage is the registration of the stitched cell membrane/wall staining image and spatial gene expression map. Since the cell nuclei and cell membrane/wall staining images have been registered in the first stage, the same operations in the second stage, include scaling, rotating, flipping and translating, are applied to the cell membrane/wall staining image (Fig. 2E). Then, the cell membrane/wall staining image and spatial gene expression map can be registered. Moreover, when utilizing staining images produced with a multi-channel microscope, STCellbin can omit the registration among these images. STCellbin can also process the case of multiple mIF staining images taken from identical tissues using the same microscope when there is only a difference in offsets among these images.

**Figure 2.**
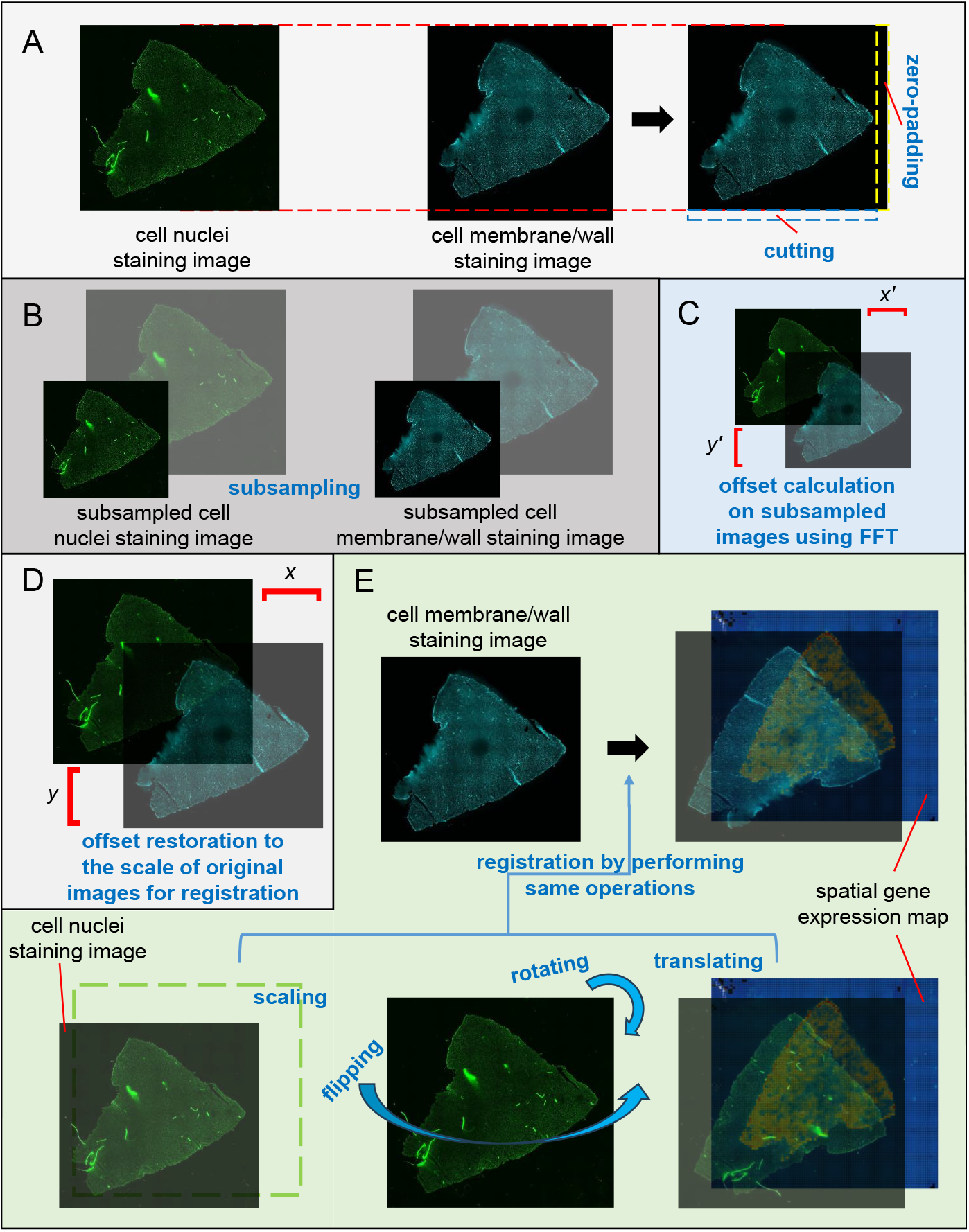
Registration of the cell membrane/wall staining image and spatial gene expression map using the cell nuclei staining image as a bridge. **A**. Size of the cell membrane/wall staining image is adjusted to be consistent with the cell nuclei staining image. **B**. Cell nuclei and cell membrane/wall staining images are subsampled. **C**. Calculating the offsets of the subsampled images. **D**. Restoring the offsets to the scale of original images for registration. E. Registering the spatial gene expression map and cell nuclei staining image by performing scaling, rotating, flipping and translating, and registering the spatial gene expression map and cell membrane/wall staining image by performing the same operations.

### Cell segmentation

The cell segmentation step of STCellbin is performed using Cellpose 2.0 [12] with some adjustments. The model architecture of Cellpose 2.0 and its weight files “cyto2” are downloaded. Due to the large size of staining images derived from Stereo-seq data, Cellpose 2.0 cannot be executed smoothly using normal hardware configurations. To circumvent this issue, the staining images are therefore cropped into multiple tiles with overlapping areas to perform cell segmentation and record the coordinates of tiles. The overlapping areas rescue cells at the border of the tiles from being cropped. To obtain the best results, segmentations with different values of the cell diameter parameter are performed independently, and the result with the largest sum of cell areas is retained. All the segmented tiles are assembled into the final segmented result according to the recorded coordinates. Moreover, when selecting the tissue segmentation option, an additional step is executed to apply a filter on the cell mask using the tissue mask, resulting in a filtered segmented outcome.

### Molecule labeling

The molecule labeling of STCellbin is the same as the one used in StereoCell in principle. StereoCell assigns molecules in the cell nuclei to the cell by using the cell nuclei mask, and then assigns molecules outside the cell nuclei to the cells with the highest probability density using Gaussian Mixture Model [15]. STCellbin assigns molecules to the cells directly based on the cell mask, while the process of assigning molecules outside the cell is included as an option. The latter decision was made as the cell membranes/walls are usually tightly packed, with only a few molecules outside the cells, and the assignment of these molecules takes a lot of time. Thus, we generally do not recommend this option, and the users can use it according to actual requirements.

## RESULTS

### Datasets and computing resource

We adopt two datasets acquired via Stereo-seq technology [4]. One is a mouse liver dataset, a tissue that offers cell boundary information via cell membranes, as in all mammalian tissues. The other dataset is derived from seeds of the plant *Arabidopsis*, a tissue that provides cell boundary information based on rigid cell walls. More details of the two datasets are shown in Table 1.

**Table 1.** Details of two datasets used for evaluation of cell boundary information.

| Detail | Mouse liver dataset | *Arabidopsis* seed dataset |
| --- | --- | --- |
| Data source | A slice of liver | Slices of multiple seeds |
| Cell nuclei dye | DAPI | ssDNA |
| Cell membrane/wall dye | mIF | CFW |
| Number of molecules | 16,177,288 | 62,884,637 |

The experiment for image segmentation is implemented on the STOmics cloud platform [16] with settings of 32 CPUs, 32 GB memory and “ALL” resource type. Except for the watershed method [17], which is implemented using ImageJ on a computer with a 16-core CPU and 16GB of RAM. The experiment for downstream analysis is implemented on a server with 40-core CPU, 128 GB of RAM and 24 GB of GPU.

### Evaluation criteria for cell segmentation performance

In a cell mask image, the gray value of a pixel is set to 255 in the cell area and 0 in the background. True positive (TP, the number of pixels with gray value of 255 in both ground truth and segmented result), true negative (TN, the number of pixels with gray value of 0 in both ground truth and segmented result), false positive (FP, the number of pixels with gray value of 0 in ground truth and 255 in segmented result) and false negative (FN, the number of pixels with gray value of 255 in ground truth and 0 in segmented result) are calculated. The number of cells segmented by a method is *ns*. For each segmented cell (cell_*i*_), there should be a corresponding area in ground truth (area_*i*_), where *i* is the cell index (*i* = 1, 2, …, *ns*). The intersection over union metric (*IoU*) [18] is set as:

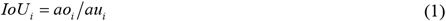

where *ao*_*i*_ is the overlap area between cell_*i*_ and area_*i*_, and *au*_*i*_ is the union area of cell_*i*_ and area_*i*_.

Then the precision (*Pre*), recall (*Rec*), F1 score (*F*1_*s*), Dice coefficient (*Dc*) and average Jaccard index (*Avg*_*J*) are calculated as:

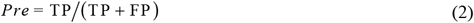

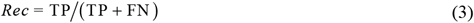

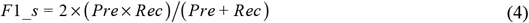

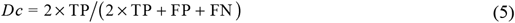

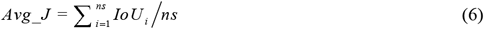

### Process and evaluation of downstream analysis

The generated single-cell spatial gene expression profiles are input into Stereopy (v0.6.0) [19]. The cells with fewer than 10 expressed genes, fewer than 3 expression counts and more than 3% mitochondrial genes are removed, and genes present in less than 3 cells are also removed. After normalization, the differentially expressed genes are summarized by Principal Component Analysis to reduce the data dimensionality, in which the number of features after reducing is 10. The Leiden algorithm [20] is used for clustering and the Uniform Manifold Approximation and Projection (UMAP) algorithm [21] is used to obtain 2D data projection. The Silhouette coefficient (*Sc*) and Moran’s I (*MI*) are used to evaluate the effect of clustering and the spatial self-correlation of each cluster, respectively. *Sc* is calculated as:

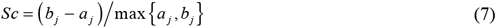

where *a*_*j*_ is the average distance between the *j*-th sample and other samples in its cluster, and *b*_*j*_ is the average distance between the *j*-th sample and the samples in other clusters. *MI* is calculated as:

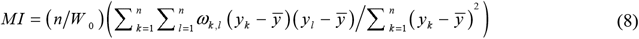

where *n* is the number of clusters, *y*_*k*_ and *y*_*l*_ are the attribute value of the *k*-th and *l*-th clusters, respectively, *y*□ is the mean of all cluster attributes, *ω*_*k,l*_ is the spatial weight between *k*-th and *l*-th clusters, and *W*_0_ is the aggregation of all spatial weights as:

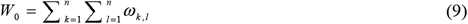

### STCellbin more accurately segments cells based on cell membrane/wall staining images

We crop two areas with higher image quality from the two datasets and design their ground truths based on the manual markup of the cells according to the cell membranes/walls in the staining images. The cell segmentation method of STCellbin is compared with the original Cellpose [18], a state-of-the-art method DeepCell [22] and a traditional watershed method [17].

On the mouse liver dataset, STCellbin effectively identifies cell membranes for segmentation, yielding cell masks that exhibit acceptable agreement with the stating image and ground truth (Fig. 3A, upper). Among all cell mask images, STCellbin provides the best description of the cell boundaries, outperforming other methods that tend to miss quite a few cells (Fig. 3A, lower). We observe a similar situation on the *Arabidopsis* seed dataset, which indicates that STCellbin can also effectively identify cell walls for segmentation (Fig. 3B). Compared with other methods, STCellbin obtains higher values in most indicators on two datasets (Fig. 3C). The comparison with the original Cellpose validates the effectiveness of STCellbin in adjusting segmentation. While DeepCell is a powerful method, it is primarily designed for segmenting cell nuclei, which involves identifying highlighted areas in the nuclei staining images. This strategy is not suitable for cell membrane/wall staining images, resulting in less desirable results. Similarly, the traditional watershed method performs poorly on cell membrane/wall staining images. In general, STCellbin’s cell segmentation is the most feasible method.

**Figure 3.**
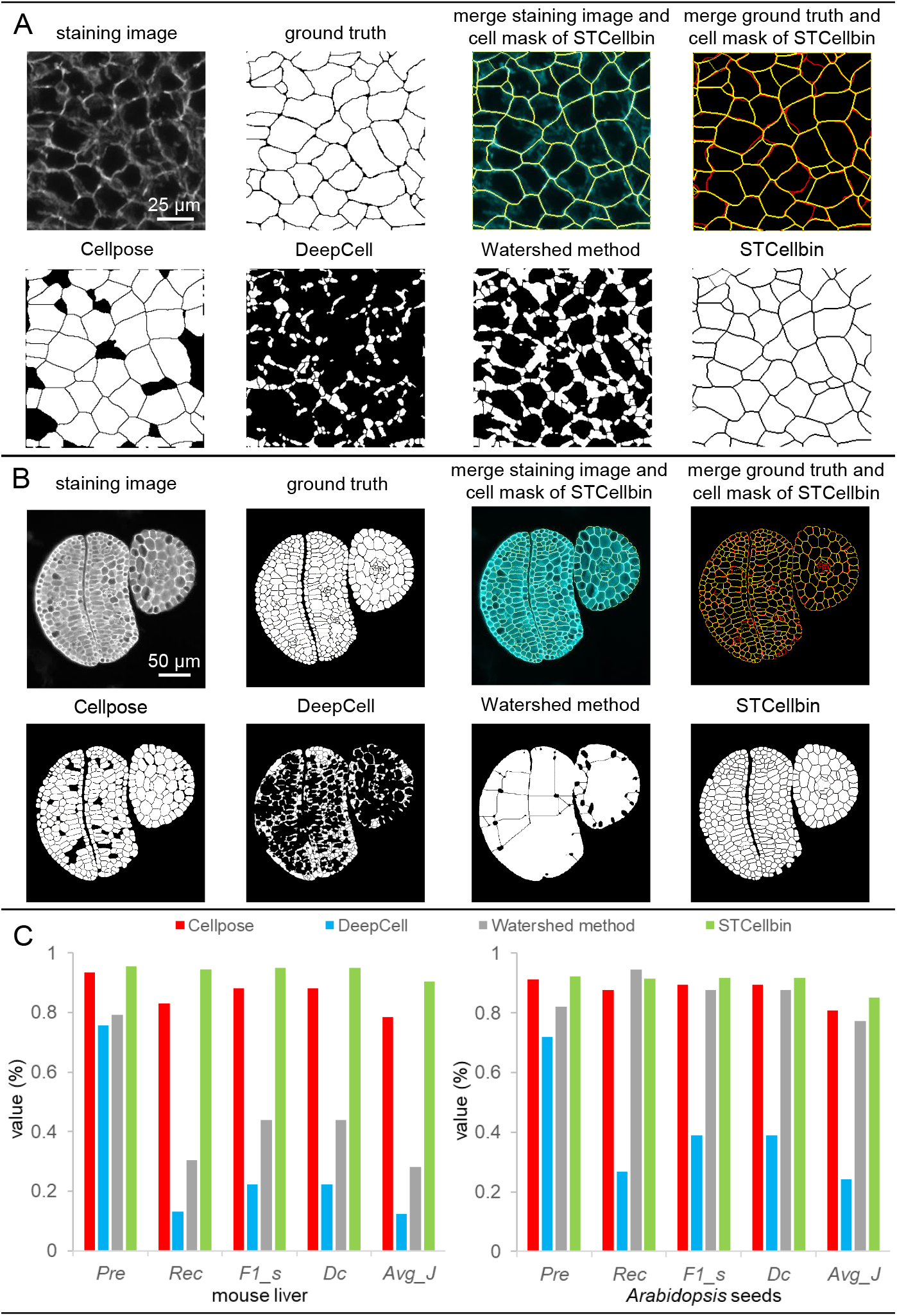
Comparison of cell segmentation performance. **A**. Cell segmentation results on the cropped area from mouse liver dataset, where in the merged images, cell masks are set in yellow, staining images are set in cyan, and ground truths are set in red. **B**. Cell segmentation results on the cropped area from *Arabidopsis* seed dataset, where in the merged images, cell masks are set in yellow, staining images are set in cyan, and ground truths are set in red. **C**. Indicator comparison of cell segmentation results on the cropped areas from mouse liver and *Arabidopsis* seed datasets.

### STCellbin generates more reliable single-cell spatial gene expression profile for downstream analysis

Currently, there is a lack of image-based one-stop software like STCellbin for Stereo-seq data. Therefore, we compare STCellbin with Baysor (v0.6.2) [23], a tool that generates the spatial gene expression profile without relying on image. However, Baysor cannot output the results on the complete mouse liver and *Arabidopsis* seed datasets in an acceptable time or a given computational resource. We run Baysor on a smaller *Arabidopsis* seed dataset, which is the cropped area in cell segmentation experiment and contains two complete seed data.

The cell area, number of unique genes per cell and number of gene counts per cell are statistically calculated from the results of STCellbin (Fig. 4A). The clustering results of STCellbin are obtained by utilizing the generated single-cell spatial gene expression profiles. The clusters of cells are spatially mapped within the tissue (Fig. 4B, left hand side for each tissue), allowing for the observation of their specific positions. From the UMAPs, it is apparent that the different cell types are effectively distinguished (Fig. 4B, right hand side for each tissue). The spatial location of the different cell types will positively influence a series of downstream analyzes such as cellular annotation in less well-studied tissues. The cells are clustered into 7 clusters on the profile of STCellbin, while 14 clusters on the profile of Baysor (Fig. 4C, the 1st subfigure from the left). We observe that the number of cells segmented by Baysor is significantly higher than that segmented by STCellbin, and it does not align with the cell count observed in the ground truth image. This may be the main reason for the higher number of clusters of Baysor. The silhouette coefficient and Moran’s I obtained by both STCellbin and Baysor are not very satisfactory (Fig. 4C, the 2nd and 3rd subfigures from the left), possibly due to the limited information from a small dataset.

**Figure 4.**
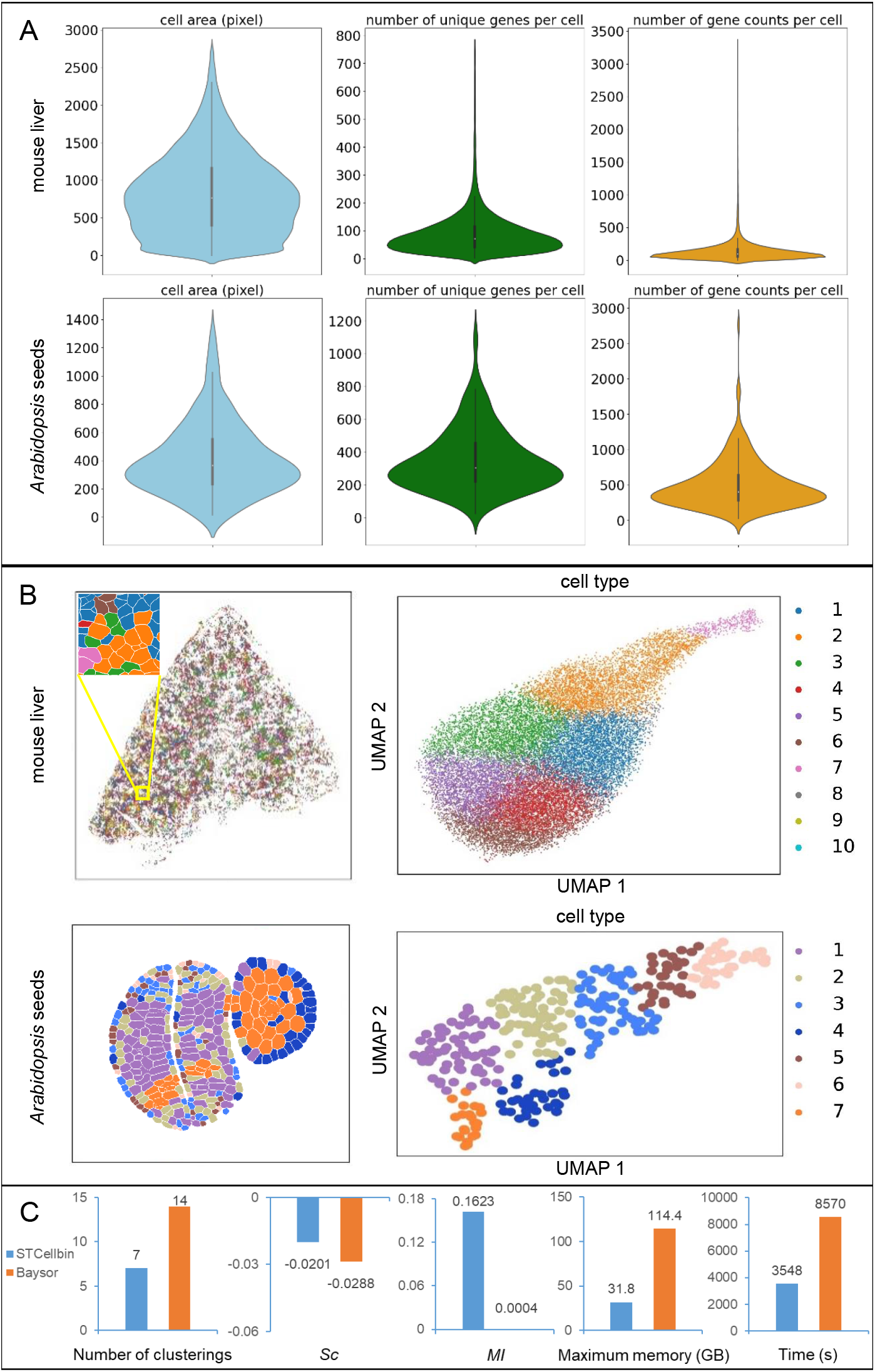
Downstream analysis results and comparison. **A**. Statistical results of cell area, gene number per cell and gene expression per cell of STCellbin on the mouse liver dataset and cropped area from *Arabidopsis* seed dataset. **B**. Clustering results and UMAPs of STCellbin on the mouse liver dataset and cropped area from *Arabidopsis* seed dataset. **C**. Indicator comparison of downstream analysis results on the cropped area from *Arabidopsis* seed dataset, where *Sc* and *MI* are taken from mean of the values of all clusters.

Nevertheless, the values of STCellbin are higher than those of Baysor. Moreover, STCellbin has significant advantages in terms of computing resource usage and running time (Fig. 4C, the 4th and 5th subfigures from the left), which also explains why Baysor cannot process the complete mouse liver and Arabidopsis seed datasets. It should be noted that the stitching and registration steps of STCellbin cannot be performed on the cropped dataset, and then, the related computational resource usage and running time cannot be recorded. Thus, the resource usage and time of STCellbin for comparison are obtained on the complete *Arabidopsis* seed dataset. That is, STCellbin can process a larger dataset with fewer computational resources and less time compared to Baysor. Overall, STCellbin is a more reliable method, particularly for analyzing high-resolution and large-field-of-view spatial transcriptomic data.

## Discussion

Accurate identification of cell boundaries plays a crucial role in generating single-cell resolution in spatial omics applications. Based on previous work in StereoCell using cell nuclei staining images to generate single-cell spatial gene expression profiles, this STCellbin update extends the capability to automatically process Stereo-seq cell membrane/wall staining images for identification of cell boundaries that facilitates downstream analyses. We also showcase a few examples of the performance of cell membrane/wall segmentation in STCellbin. Currently, the tools for cell nuclei and cell membrane/wall segmentation can be independently executed, allowing users to choose the more suitable solution for their specific applications. In future work, these two techniques can be combined by training a deep learning model that is compatible with any staining image type, thereby achieving more accurate results.

## AVAILABILITY OF SOURCE CODE AND REQUIREMENTS

- Project name: STCellbin
- Project home page: https://github.com/STOmics/STCellbin
- Operating system(s): Platform independent
- Programming language: Python
- Other requirements: Python 3.8
- License: MIT License
- RRID: SCR_024438

## DATA AVAILABILITY

The data that support the findings of this study have been deposited into Spatial Transcript Omics DataBase (STOmics DB) of China National GeneBank DataBase (CNGBdb) with accession number STT0000048: https://db.cngb.org/stomics/project/STT0000048. A backup link for the data is also provided at Github link of STCellbin: https://github.com/STOmics/STCellbin.

## LIST OF ABBREVIATIONS

DAPI: 4,6-diamidino-2-phenylindole
H&E: hematoxylin-eosin
ssDNA: single strand DNA fluorescence
mIF: multiplex immunofluorescence
CFW: calcofluor white
FFT: Fast Fourier Transform.

## DECLARATIONS

### Ethics Approval and Consent to Participate

Not applicable.

### Competing Interests

The authors declare that they have no competing interests.

### Funding

This work was supported by the National Key R&D Program of China (2022YFC3400400).

### Authors’ Contributions

Conceptualization: BZ and ML; Project administration and supervision: SB and XX; Software implementation: ZD, HQ, KS and HL; Data collection and processing: QK, XF and LC; Validation: QK and ZD. Project coordination: BZ and ML; Manuscript writing and figure generation: BZ, ML and QK; Manuscript review: ML, SF, YZ, YL and SB.

## Acknowledgements

We thank China National GeneBank for providing technical support.

## Notes

### Competing Interest Statement

The authors have declared no competing interest.

### Summary of Updates

We have added new experiments and corrected grammatical and writting errors in the paper.

